# Sub-zero temperatures during early spring migration in blue-spotted salamanders (*Ambystoma laterale*)

**DOI:** 10.1101/2025.03.28.645960

**Authors:** Danilo Giacometti, Patrick D. Moldowan, Glenn J. Tattersall

## Abstract

Amphibians that reproduce in early spring at northern latitudes may encounter environmental ice while migrating to their breeding sites. Due to the nucleation properties of ice, contact with environmental ice may induce rapid freezing of body tissues, which can cause irreversible damage to cells and lead to death. Although some species of salamanders are known to move over ice during early spring migration, freeze-intolerant species are expected to avoid physical contact with ice crystals to minimise the risk of freezing. Here, we documented the thermal biology of the freeze-intolerant blue-spotted salamander (*Ambystoma laterale* Hallowell, 1856) migrating at sub-zero temperatures in Algonquin Provincial Park, Ontario, Canada. During our surveys, we found sheltered, inactive, and migrating individuals; some in direct contact with ice. Our field measurements of skin temperature using high resolution thermal imaging suggest that *A. laterale* can sustain activity in a supercooled state (i.e., chilled below the freezing point of body fluids but not frozen). By migrating in a supercooled state, these salamanders may overcome the risk of freezing while simultaneously prolonging their breeding season and potentially avoiding predators.

## Introduction

In spring, northern populations of *Ambystoma* salamanders emerge from underground burrows and migrate overland to breeding ponds after months of fasting and prolonged exposure to cold (Wright and Allen 1909; Smallwood 1928). The blue-spotted salamander (*Ambystoma laterale* Hallowell, 1856) (**Fig. 1a**) is a freeze-intolerant amphibian known to migrate in late winter and early spring, when sub-zero temperatures are frequent, and ice and snow cover persist (Storey and Storey 1986; Lowcock et al. 1991). Thus, it is possible that salamanders risk freezing while partaking in breeding migration (Madison 1997).

**Fig 1.**
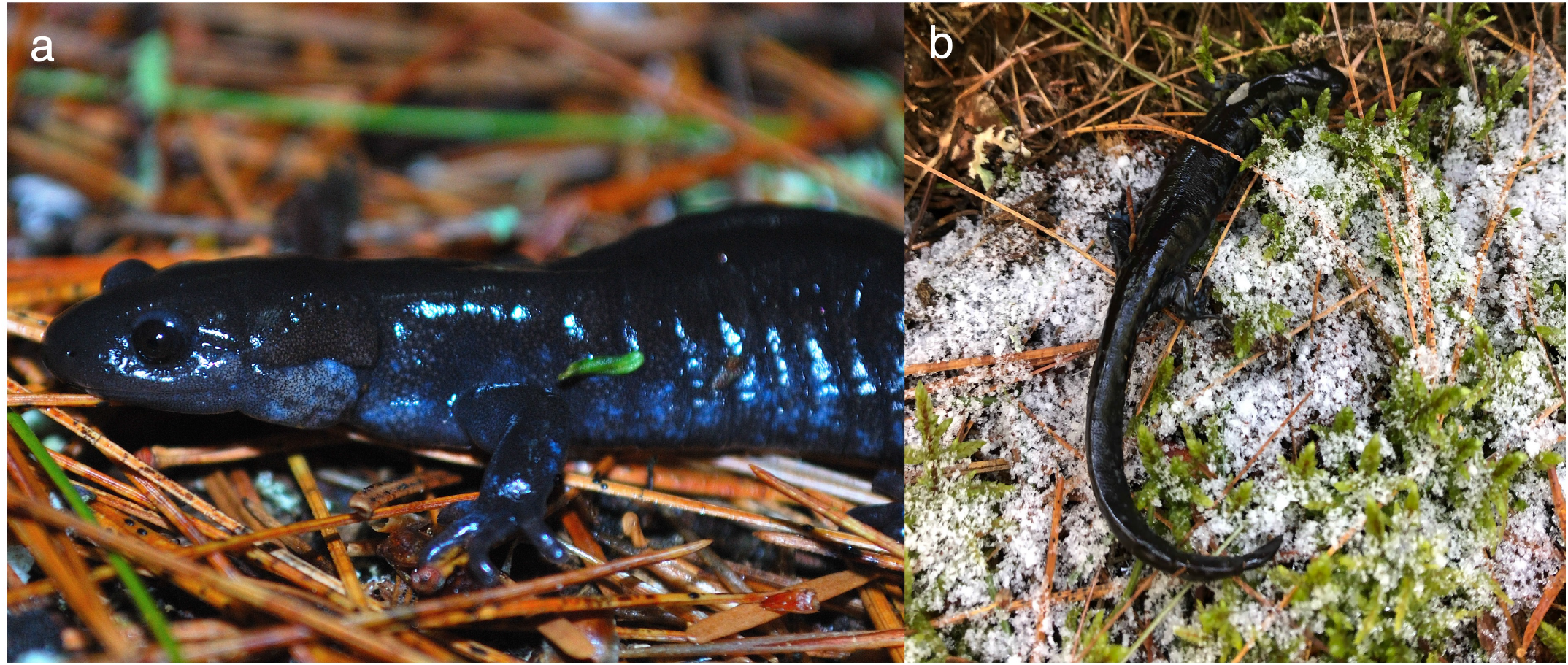
**a**. Male *Ambystoma laterale* Hallowell, 1856 over needles of Eastern White Pine (*Pinus strobus* L., 1753) on the forest floor surrounding Bat Lake, Algonquin Provincial Park, Ontario, Canada. **b**. Salamander observed in direct contact with ice crystals. Photos by Danilo Giacometti.

The negative effects of freezing are myriad, ranging from irreparable cell damage to death (Storey and Storey 1992). The freezing point of most vertebrate tissues in the absence of cellular antifreezes (∼ –0.5 °C) is similar to that of water (Costanzo et al. 1995). Laboratory evidence suggests that *A. laterale* are capable of supercooling to –1.5 ºC (i.e., chilling below the freezing point of their body fluids without freezing) (Storey and Storey 1986). However, this strategy cannot be sustained for longer than 24 h, because, as amphibians, salamanders possess permeable skin that does not prevent environmental ice from entering their bodies and inducing freezing (Storey and Storey 1986).

Although some species of salamanders are known to move over ice (e.g., Blanchard 1930; Catenazzi 2016), we still have limited information about salamander thermal biology in natural settings, especially during breeding migration (Giacometti et al. 2024). Given its intolerance to freezing, one could expect that *A. laterale* should behaviourally avoid physical contact with environmental ice to reduce exposure to external nucleators (Lee and Costanzo 1998). In doing so, the salamanders would minimise the risk of freezing, maximise opportunity for overland movement, and increase their odds of a successful breeding season. Here, we show that field-active *A. laterale* migrate at temperatures below their known minimum supercooling point even while in contact with ice crystals (**Fig. 1b**).

## Material and methods

### Salamander sampling

On 22 and 23 April 2022, we studied a population of *A. laterale* at Bat Lake, Algonquin Provincial Park, Ontario, Canada (45.5857ºN, 78.5185ºW) with authorisation from Brock University’s Animal Care Committee (AUP #22-03-04), the Ministry of Northern Development, Mines, Natural Resources and Forestry (#1100575), and Ontario Parks. Bat Lake is acidic (Ph ∼4.5–4.8), fishless, and has permanent water (Bianchini et al. 2012; Moldowan et al. 2022). This population of *A. laterale* co-occurs with a low frequency of *A. laterale* (2)-*jeffersonianum* (LLJ) genotype (∼7.6% LLJ) (COSEWIC 2016); we did not distinguish between the two genotypes in the present study. In 2017, a drift fence was installed around the perimeter of Bat Lake for an amphibian population study. Logs and wood boards were placed along the fence to serve as refuges for amphibians that reach the fence but are unable to cross it. Nighttime surveys are conducted throughout the spring migratory period to ensure the timely processing and movement of amphibians (Moldowan 2023).

To search for inbound, pre-breeding *A. laterale*, we conducted visual surveys and flipped over the logs and wood board hides along the drift fence. We took a thermal image of encountered salamanders using a FLIR SC660 thermal camera (FLIR Systems). This camera has a resolution of 640 x 480 pixels, functions between a temperature range of –40 to 120ºC, and has a measurement accuracy of ±1 ºC (Playà_JMontmany and Tattersall 2021). At the site of sampling, we also measured air temperature (*T*_air_) and relative humidity at ∼1.50m above ground using a Kestrel 4000 (Kestrel Meters™). After imaging an individual, we sexed it based on secondary sexual characteristics. Specifically, we visually inspected cloacal morphology to assess whether the individual had pleated (female) or papillose (male) cloacae (Petranka 1998). We also recorded the body mass (M_b_) of each imaged individual with a Pesola™ spring scale (to the nearest 0.1g). After body size measuring, we returned all individuals to their original site of sampling.

### Thermal imaging processing

We processed all thermal images using ThermimageJ plugins for Fiji that were verified against algorithms in FLIR Research Studio Software (Tattersall 2019). For all images, we assumed an object emissivity of 0.95 in line with previous work investigating the thermal biology of amphibians through thermography (Rowley and Alford 2007). To ensure proper extraction of temperature data from our thermographs, we verified camera calibration constants and adjusted object parameters (e.g., object distance, relative humidity, atmospheric temperature, reflected temperature) according to conditions measured at the site of collection. To obtain measurements of skin temperature (*T*_skin_), we digitally drew a region of interest (ROI) over the dorsum of the animal and recorded the average temperature. We also drew an oval ROI of the same size over the ground immediately adjacent to the animal to estimate an average ground temperature (*T*_ground_). From each ROI, we obtained an average temperature and its associated standard deviation; standard deviation was low for all measurements, indicating consistency in our measurements. We assumed that *T*_skin_ would match core body temperatures primarily due to the small size of *A. laterale*, but also based on preliminary evidence suggesting a strong correlation between *T*_skin_ and core temperatures in frogs (Rowley and Alford 2007); data from lizards of comparable M_b_ to our study system also support this observation (Barroso et al. 2016).

### Statistical analysis

Using R (version 4.3.2) in RStudio (version 2024.04.0) (R Core Team 2024), we assessed the relationship between *T*_skin_ and *T*_ground_ through Spearman’s rank correlation coefficient (ρ) implemented with the *cor*.*test* function (stats package) (R Core Team 2024). To this end, we set the parameters method = ‘spearman’ and exact = TRUE. We created a figure with the “ggplot2” package (Wickham 2016).

## Results and discussion

We imaged a total of 21 *A. laterale* (13 males and 7 females) during our surveys, which included individuals moving out in the open (migrating), stationary and out in open (inactive), and sheltered under fence-adjacent hides (sheltered). Females were heavier than males (average ± standard deviation; M_b_ females = 6.37 ± 1.66 g and M_b_ males = 4.11 ± 0.79 g). It did not rain on the nights we conducted our study. Relative humidity ranged from 64.9% to 90% (average 82.0 ± 7.61%), while *T*_air_ ranged from 0.4ºC to 4.4ºC (average = 2.89 ± 1.24ºC), and *T*_ground_ ranged from –3.28ºC to 0.11ºC (average = –1.18 ± 0.98 ºC). *T*_skin_ ranged from –3.59ºC to 0.49ºC (average = –1.07 ± 1.19 ºC), and these values are similar to those reported for supercooled *A. laterale* under laboratory conditions (Storey and Storey 1986).

We imaged six individuals moving toward Bat Lake (**Fig. 2a**). Their average *T*_skin_ was – 2.63 ± 0.75ºC, which suggests that these salamanders were sustaining activity in a supercooled state. Additionally, we found other five salamanders wedged between logs and the frozen ground (average *T*_skin_ = –0.24 ± 0.65ºC, range –1.08ºC–0.49ºC; average *T*_ground_ = –1.03 ± 0.24ºC, range – 1.38ºC–-0.76ºC). These individuals were in direct contact with ice crystals, showing that freeze-intolerant amphibians do not necessarily avoid physical contact with environmental ice to prevent freezing (Lee and Costanzo 1998). In one situation we observed two salamanders nearly adjacent to one another, with one individual almost 2ºC warmer than the other, suggesting either that these individuals were migrating rapidly from different thermal microhabitats or that one individual was undergoing freezing and releasing heat of fusion in the process (**Fig. 2b**).

**Fig. 2.**
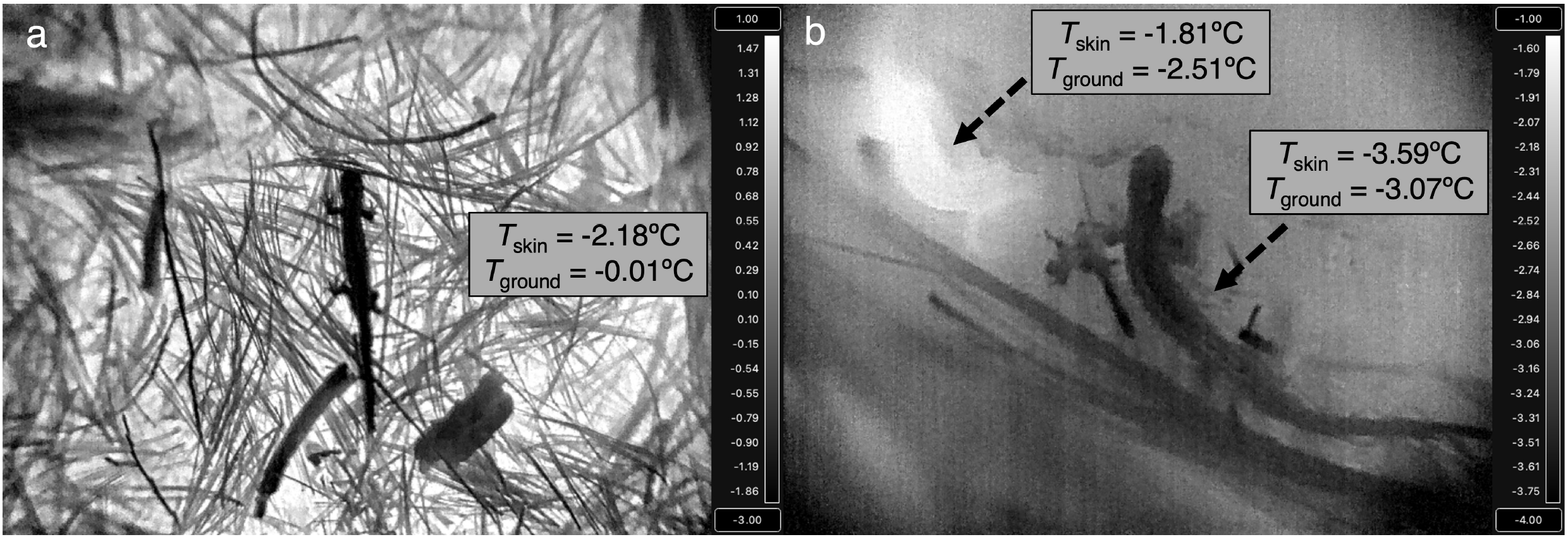
**a**. Thermal image of an *Ambystoma laterale* Hallowell, 1856 migrating at sub-zero temperatures. The temperature difference between salamander skin (*T*_skin_) and the ground (*T*_ground_) suggests that this individual was moving from a cooler substrate to a warmer site in a supercooled state. **b**. Thermal image of two nearly adjacent salamanders with contrasting *T*_skin_ standing on ice-covered ground.

It is well known that ectotherm body temperatures correlate with microhabitat temperature (Angilletta 2009). Our results support this pattern, as evidenced by the positive correlation between *T*_skin_ and *T*_ground_ (ρ = 0.50, *p* = 0.02) (**Fig. 3**). One individual, however, showed a substantially lower *T*_skin_ than what it would be predicted based on its corresponding *T*_ground_, suggesting it had not yet equilibrated close to local *T*_ground_. This lends further support to the idea that *A. laterale*, at least partially, overcome the risk of freezing by migrating while in a supercooled condition. Importantly, however, while we assumed that *T*_skin_ matched core body temperatures, the relationship between these two variables is not well understood in amphibians, particularly salamanders (Rowley and Alford 2007; Giacometti and Tattersall 2024).

**Fig. 3.**
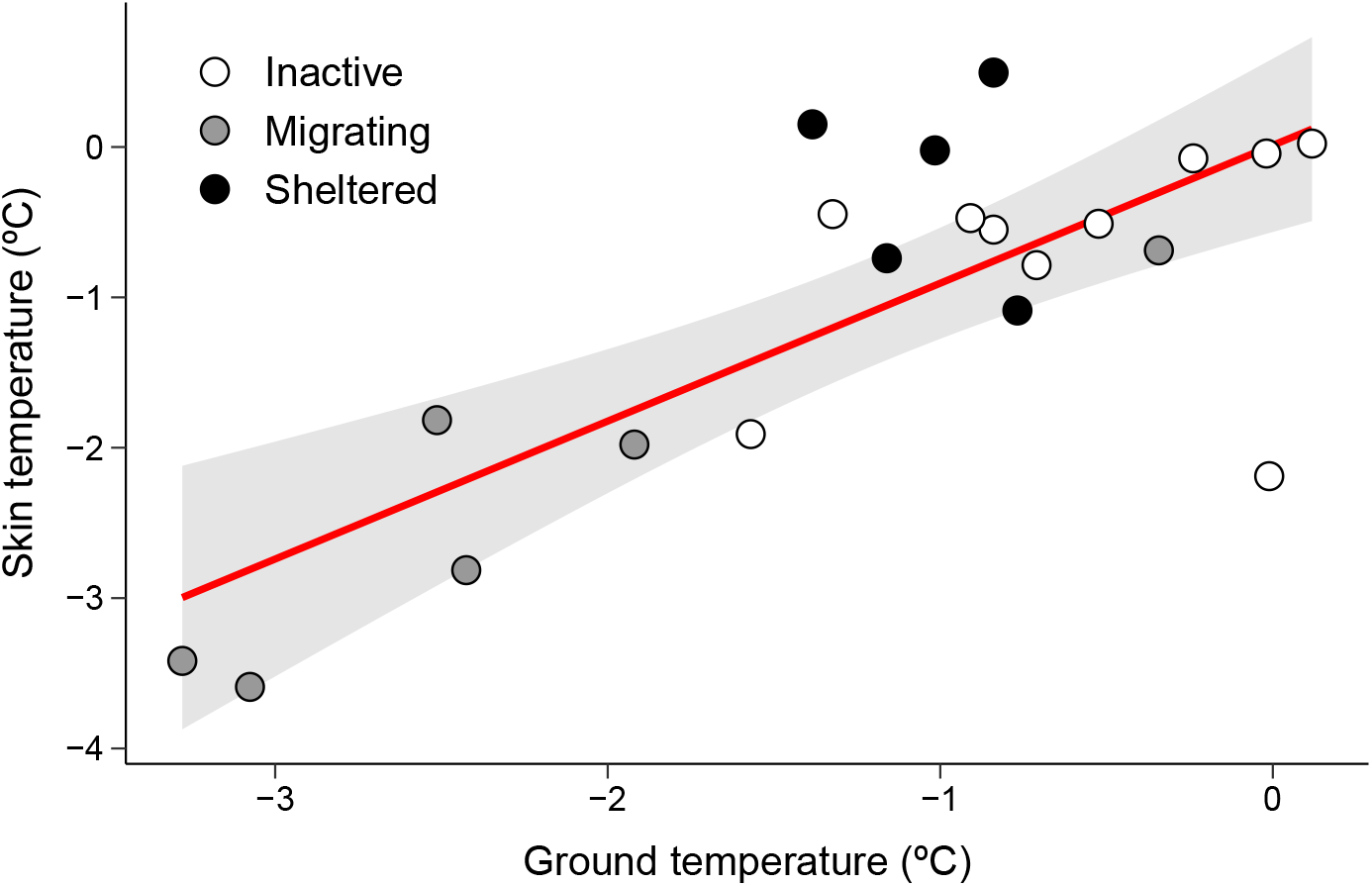
Correlation between skin temperature and ground temperature in *Ambystoma laterale* Hallowell, 1856. Each dot represents an individual salamander. The solid line and shaded area indicate the predicted relationship between skin and ground temperatures, and the 95% confidence interval, respectively.

Although freeze-intolerant, *A. laterale* have been shown to accumulate glucose—an effective cryoprotectant—in blood and liver (Storey and Storey 1986). While our observational data cannot clarify if salamanders from the Bat Lake population can withstand prolonged exposure to sub-zero temperatures, their ability to sustain activity in a supercooled state should be an important asset in their toolkit for survival. Indeed, by starting their breeding migration in late winter and early spring, *A. laterale* are the first species of amphibians to arrive at Bat Lake (Moldowan 2023). Thus, by migrating at sub-zero temperatures, *A. laterale* may limit temporal overlap with other spring breeders, prolong their breeding season, and minimise their odds of being found by predators.

## Author Contributions

Conceptualization, Writing – Original Draft Preparation: D.G., P.D.M., and G.J.T.; Writing – Review & Editing: D.G., P.D.M., and G.J.T.; Investigation: D.G. and G.J.T.; Data Curation: D.G. and G.J.T.; Formal Analysis: DG; Visualization: D.G. and G.J.T.; Supervision: G.J.T.; Funding Acquisition: G.J.T.

## Data availability statement

The full set of thermal images and data we used in the study can be accessed from Brock University Dataverse at: https://doi.org/10.5683/SP3/FZJBQH.

## Acknowledgements

We thank Ontario Parks, the Ministry of Northern Development, Mines, Natural Resources and Forestry, and the Algonquin Wildlife Research Station for facilitating access to our field site and study animals.

## Funding statement

Research funding was provided by a Natural Sciences and Engineering Research Council of Canada Discovery Grant to G.J.T. (RGPIN-2020-05089) and by Brock University through a Gervan Fearon Graduate Scholarship awarded to D.G.

## Competing interests

None declared.

## Notes

https://doi.org/10.5683/SP3/FZJBQH

### Competing Interest Statement

The authors have declared no competing interest.

